# Misspelled-word reading modulates late cortical dynamics

**DOI:** 10.1101/2025.01.05.631357

**Authors:** Jiaxin You, Aino Saranpää, Tiina Lindh-Knuutila, Marijn van Vliet, Riitta Salmelin

**Affiliations:** Department of Neuroscience and Biomedical Engineering, Aalto University, P.O. Box 12200, FI-00076, Aalto, Finland; Aalto NeuroImaging, Aalto University, P.O. Box 12200, Aalto, FI-00076, Finland

## Abstract

Literate humans can effortlessly interpret tens of thousands of words, even when the words are sometimes written incorrectly. This phenomenon suggests a flexible nature of reading that can endure a certain amount of noise. In this study, we investigated where and when brain responses diverged for conditions where misspelled words were resolved as real words or not. We used magnetoencephalography (MEG) to track the cortical activity as the participants read words with different degrees of misspelling that were perceived to range from real words to complete pseudowords, as confirmed by their behavioral responses. In particular, we were interested in how lexical information survives (or not) along the uncertainty spectrum, and how the corresponding brain activation patterns evolve spatiotemporally. We identified three brain regions that were notably modulated by misspellings: left ventral occipitotemporal cortex (vOT), superior temporal cortex (ST), and precentral cortex (pC). This suggests that resolving misspelled words into stored concepts involves an interplay between orthographic, semantic, and phonological processing. Temporally, these regions showed fairly late and sustained responses selectively to misspelled words. Specifically, an increasing level of misspelling increased the response in ST from 300 ms after stimulus onset; a functionally fairly similar but weaker effect was observed in pC. In vOT, misspelled words were sharply distinguished from real words notably later, after 700 ms. A linear mixed effects (LME) analysis further showed that pronounced and long-lasting misspelling effects appeared first in ST and then in pC, with shorter-lasting activation also observed in vOT. We conclude that reading misspelled words engages brain areas typically associated with language processing, but in a manner that cannot be interpreted merely as a rapid feedforward mechanism. Instead, feedback interactions likely contribute to the late effects observed during misspelled-word reading.

## 1 Introduction

Visual word recognition involves initial visual perception, followed by subsequent orthographic, phonological, and semantic processing (Grainger, 2008). This recognition process is effortlessly completed for familiar words because the perceived visual information matches the orthography (i.e., spelling) of an entry in the mental lexicon (Martin, Tan, Newsome, & Vu, 2017). However, when a word is misspelled, it is technically a nonword with no entry in the mental lexicon. Yet, we can often still readily identify them even if the orthography is not perfectly matched (Davis, 2013; Pearson, Barr, Kamil, & Mosenthal, 1984). Misspelled words, which are graphemically related to the real words, are not retained in long-term memory, but still plausibly involve the lexical system, potentially aiding in the retrieval of analogous words (Almeida & Poeppel, 2013). This reading flexibility effect has its counterpart in the “Ganong effect” in speech perception where listeners tend to identify an ambiguous sound as part of a real word instead of a nonword (Gow Jr, Segawa, Ahlfors, & Lin, 2008).

In masked priming, a misspelled word created by transposing two medial letters (TL) or replacing a medial letter by a different one (RL) results in a strong priming effect on lexical decision responses to a subsequent correctly spelled target word (Forster, Davis, Schoknecht, & Carter, 1987). This priming effect is in line with the repetition account, which states that misspelled words access the lexical entries for the correctly spelled ones (Forster et al., 1987). However, when word forms are sufficiently distorted they become unrecognizable. As the number of replaced letters in the primes increases, priming effects decrease and eventually vanish (Grainger, 2008; Lupker & Davis, 2009). The absence of priming effects indicates that the severely misspelled-word primes fail to access the lexical entries of correctly spelled-word targets. Evidently, there is no strict dichotomy between words and nonwords, but rather a spectrum, with misspelled words constituting a gray area (Chen, Davis, Pulvermüller, & Hauk, 2015; Hauk, Pulvermüller, Ford, Marslen-Wilson, & Davis, 2009). While the behavioral results provide a measurement at the end of target word recognition, it remains to be clarified how the recognition of misspelled words proceeds in our brain in space and time.

When we encounter unfamiliar words of our native language, an everyday phenomenon during language development is that those words initially appear as pronounceable non-words, i.e., pseudowords. Although devoid of meaning, pseudowords are postulated to elicit a broad search in the mental lexicon (Grainger & Jacobs, 1996). In this sense, reading pseudowords might serve as a good entry point for conceptualizing the processing of mis-spelled words (Grainger & Jacobs, 1996). Functional magnetic resonance imaging (fMRI) studies have compared neural activation patterns evoked by word and pseudoword reading. Stronger activation to pseudowords than real words has often been reported, particularly in the left ventral occipito-temporal cortex (vOT) and left frontal operculum(Carreiras, Mechelli, Estévez, & Price, 2007; Cattinelli, Borghese, Gallucci, & Paulesu, 2013; Fiebach, Friederici, Müller, & Von Cramon, 2002; Heim et al., 2005; Kronbichler et al., 2007; McNorgan, Chabal, O’Young, Lukic, & Booth, 2015; Taylor, Rastle, & Davis, 2013). These findings may provide clues about where the misspelled word recognition occurs.

EEG has been used to investigate indirectly the time course of processing misspelled words by applying the masked priming paradigm, similar to previous behavioral studies. In those EEG studies, the misspelled words, while orthographically similar to words, were not always pronounceable. The priming effects were mostly reflected in the eventrelated potential (ERP) components N250 and N400, negative-going responses that reach the maximum at around 250 ms and 400 ms post stimulus onset, respectively. Specifically, a nonword prime with a replaced letter that was visually dissimilar to the target word (e.g., dentgst-DENTIST) resulted in stronger N250 and N400 responses to the target word than when the replaced letter was visually similar to the target (e.g., dentjst-DENTIST) (Gutíerrez-Sigut, Marcet, & Perea, 2019). Enhancement of N250 and N400 for the target words was also observed with primes that contained double replacement (e.g., shgue-SHAPE) compared to primes that contained transposed letters (e.g., shpae-SHAPE) (Ktori, Kingma, Hannagan, Holcomb, & Grainger, 2014; Meade, Grainger, & Holcomb, 2021; Meade, Grainger, Midgley, Holcomb, & Emmorey, 2020; Meade, Mahnich, Holcomb, & Grainger, 2021). The different types of misspelled words used as primes thus facilitated the processing of word targets to a varying degree; however, in these masking studies, the primary focus was on word targets rather than on the misspelled words themselves. In contrast, during single-word reading, only a stronger N400 response was reported for RL misspelled (and pronounceable) words compared to TL pseudowords and real words (Vergara-Martínez, Perea, Gómez, & Swaab, 2013). Effects related to N250 thus seem to be specifically related to the masked priming paradigm. This suggests that N250 is likely linked to the sublexical orthographic overlap between prime and target, as it shows a gradient modulated by their orthographic similarity (Gutíerrez-Sigut et al., 2019). Presumably, the later stages of processing, including the N400 component, are of primary interest when studying cortical dynamics of recognition of isolated misspelled words. N400 is thought to reflect lexical and semantic processing at the whole-word level (Kutas & Federmeier, 2011). It may indicate the amount of effort required to translate a word form into its corresponding semantic concept (Holcomb, Grainger, & O’rourke, 2002). According to the latest predictive coding model, the heightened N400 response observed for pseudowords reflects an increased lexico-semantic prediction error Eddine, Brothers, Wang, Spratling, and Kuperberg (2024). It remains unclear whether this framework can be applied to understand the neural dynamics involved in reading misspelled words.

Magnetoencephalography (MEG) has become an increasingly prevalent method to study language function in the brain due to its combined temporal and spatial sensitivity (Salmelin, Kujala, & Liljeström, 2019). MEG has revealed a salient spatiotemporal process for single-word reading that proceeds from analysis of visual features at around 100 ms in the occipital cortex, to letter-string processing at around 150 ms in the left occipitotemporal cortex, and ultimately to lexical and semantic processing at around 200– 800 ms in the left superior temporal cortex (Tarkiainen, Helenius, Hansen, Cornelissen, & Salmelin, 1999; Vartiainen, Liljeström, Koskinen, Renvall, & Salmelin, 2011). The N400 (or N400m for MEG) response in the left superior temporal cortex is stronger and longer lasting for pseudowords than real words (Vartiainen et al., 2011; Wydell, Vuorinen, Helenius, & Salmelin, 2003). However, no effects were observed in the left occipitotemporal cortex, despite its sensitivity to pseudowords in previous fMRI studies (Woolnough et al., 2022, 2021). This discrepancy relights the debate regarding the functional role of the left occipitotemporal cortex—whether it primarily supports prelexical processing (Baker et al., 2007; Dehaene, Le Clec’H, Poline, Le Bihan, & Cohen, 2002; Glezer, Jiang, & Riesenhuber, 2009; Tarkiainen et al., 1999; Vartiainen et al., 2011) or engages in top-down lexical or phonological information (Dehaene & Cohen, 2011; Price & Devlin, 2011; Woolnough et al., 2021). The modulation of cortical response by misspelled words remains to be investigated, which may offer new insights into this debate.

The present study aimed to determine whether these previously described neural effects of single-word reading serve as indicators for misspelled word resolution and whether they might also be linked to the degree of word-likeness along the spectrum from word to pseudoword. Starting from base words, we parametrically replaced an increasing number of letters while keeping the misspelled words pronounceable. In previous behavioral studies, letter replacement has been used to effectively manipulate misspelling. This approach provided a continuum of misspelled words, ranging from real words to complete pseudowords. MEG and concurrent behavioral data were recorded in a visual word recognition paradigm.

## 2 Materials and Methods

### 2.1 Participants

Twenty-five volunteers participated in this study (19 females and 6 males; age range 19–40 years; mean age 24.4 years and SD 5.5). All participants were healthy, right-handed native Finnish speakers, without history of language disorders or problems in language development, or psychiatric, neurological or somatic disorders. A written informed consent was obtained from all participants, in accordance with the prior approval of the Aalto University Research Ethics Committee. Data of two subjects were discarded from the analysis: one due to excessive noise from head movements, and the other due to non-compliance with the task as evidenced by both self-report and behavioral results. Consequently, data from 23 participants were retained for analysis in this study.

### 2.2 Stimuli

Four stimulus categories, in increasing order of orthographic dissimilarity, were real Finnish words (RW), and three levels of misspelled words constructed by replacing 1, 2, or 3 internal letters (vowel with vowel and consonant with consonant) of the original base words, respectively (RL1, RL2 and RL3) (Figure 1b). In this manner, the stimuli were intended to exhibit graded levels of recognition difficulty. Each category contained 150 stimuli, and each stimulus was derived from a different base word to avoid possible word-form priming effects between stimuli. All 600 base words were frequent Finnish nouns with lengths of 7-8 letters, selected from a large Finnish Internet corpus (Kanerva, Luotolahti, Laippala, & Ginter, 2014). The base words in the four categories did not differ significantly in lemma frequency (one-way ANOVA test, *p* = .81). Table 1 summarizes the psycholinguistic characteristics of each stimulus category.

**Figure 1:**
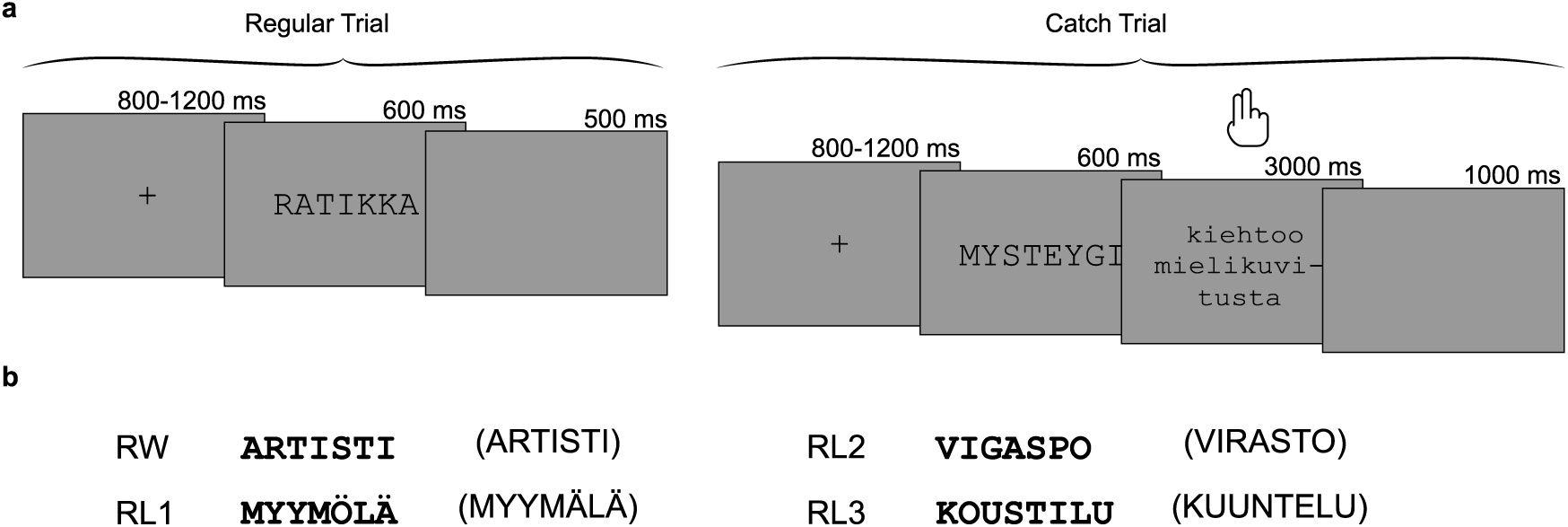
**a**, Experimental design. Of all trials, 90% constituted regular trials, included in the MEG data analysis, while the remaining 10% were catch trials. During regular trials, participants engaged in silent reading of the presented stimuli, whereas in catch trials, participants were required to make a behavioral response on sentence validity (“MYSTYGY fascinates the imagination”). **b**, Example stimuli from each of the four stimulus categories, along with their corresponding base words enclosed in parentheses (ARTISTI = artist, MYYMÄLÄ = store, VIRASTO = bureau, KUUNTELU = listening).

**Table 1:** Summary of the psycholinguistic characteristics for each stimulus category (mean±std).

| Category | #Replaced letters | Length | Frequency (log)<sup>a</sup> | Visual distance (pixel)<sup>b</sup> | Bigram frequency<sup>c</sup> |
| --- | --- | --- | --- | --- | --- |
| RW | 0 | 7.28 ± 0.45 | 11.51 ± 0.94 | 0 | 66168.83 ± 20861.75 |
| RL1 | 1 | 7.60 ± 0.49 | 11.55 ± 0.97 | 3555.73 ± 377.07 | 54467.24 ± 23649.89 |
| RL2 | 2 | 7.64 ± 0.48 | 11.48 ± 0.92 | 4938.47 ± 419.42 | 39156.36 ± 16258.14 |
| RL3 | 3 | 7.40 ± 0.49 | 11.52 ± 1.03 | 6123.21 ± 441.38 | 41067.07 ± 18433.07 |

<sup>a</sup> Frequency: Log lemma frequency value of corresponding base word.
<sup>b</sup> Visual distance: Euclidean distance between the corresponding pixels of the two rendered images of stimulus and its base word.
<sup>c</sup> Bigram frequency: averaged number of times the consecutive pairs of letters within a stimulus appear in the used corpus.

All categories of misspelled words were readable pseudowords, which tend to engage subjects in lexical inference (and implicitly more than for nonwords (Evans, Lambon Ralph, & Woollams, 2012; Schuster, Hawelka, Richlan, Ludersdorfer, & Hutzler, 2015)). The pronounceability was controlled according to Finnish phonotactics (Suomi, Toivanen, & Ylitalo, 2009) to conform to Finnish orthographic rules. Using letter replacement, it was simultaneously feasible to parametrically vary the orthographic dissimilarity (i.e., misspelling) without changing the stimulus length and retain the misspelled words pronounceable (i.e., pseudoword). This is not always possible for other letter manipulation approaches such as letter transpose, deletion, or addition.

To prevent lexical competitor effects, the generated pseudoword was ensured to be closest orthographically to its corresponding base word, i.e., any other words in the corpus should be farther away from the base word based on Damerau–Levenshtein distance (Damerau, 1964; Levenshtein, 1965). However, this requirement could not be met for RL3 as there were always other words with a Damerau–Levenshtein distance of 3 from the base word. In this sense, RL3 has no unique map to a real word, thereby remaining unresolved for lexical retrieval.

### 2.3 Experimental procedure

Stimuli were presented one by one in a pseudorandom order using Presentation software (Neurobehavioral Systems Inc., USA). Participants were asked to read the stimuli silently and attempt to retrieve their corresponding base words. We chose a silent reading task, instead of a lexical/phonological decision task, since we aimed to simulate natural reading where we typically do not make any decisions but try to understand the content while reading. In each trial, a fixation cross was presented for 800-1200 ms on the screen with a gray background. Next, a stimulus was presented in black capital letters in 36-point Courier New font for 600 ms (visual angle ≈ 0.61*^◦^*), followed by a 500-ms blank gray background.

To keep the participants engaged, 10 % of the trials in each category were designated as catch trials, during which a stimulus was followed by a sentence missing the first word. Participants were instructed to determine whether the base word inferred from the preceding stimulus could reasonably serve as the missing first word of the sentence (Hultén et al., 2021). For example, a stimulus “MYSTYGY”, an RL2 from the base word of “MYSTERY”, might be followed by “stimulates the imagination” (for the actual Finnish-language stimuli: MYSTEYGI (MYSTEERI) kiehtoo mielikuvitusta). In this case the answer would be “yes” even though the stimulus was not a correctly-written word. Participants had 3 seconds after the sentence onset to respond with a button press (Figure 1a). The responses, including reaction time and accuracy, were obtained from catch trials and used as behavioral data; the corresponding MEG trials were excluded from further analysis as, compared to the other trials, they contained additional stimuli and neural activity. To minimize fatigue, subjects were given five self-paced breaks during the experiment.

After the experiment, we asked participants to complete a brief questionnaire to evaluate how they engaged in this experiment. During this questionnaire, they wrote down self-reported feedback on a number of questions: fatigue, stimulus pronounceability, speed of stimulus presentation, percentages of recognizable and unrecognizable stimuli, strategies used to recognize misspelled words, and additional comments. The results from the questionnaire were not used in any analysis.

### 2.4 Data acquisition

MEG data were measured at the Aalto NeuroImaging (ANI) MEG Core with a MEGIN TRIUX neo system (MEGIN Oy, Helsinki, Finland). The system is equipped with 102 triplet sensors, with each triplet containing two orthogonal planar gradiometers and one magnetometer. Participants were seated in the MEG device in a magnetically shielded room (Imedco AG, Switzerland).The data was low-pass filtered at 330 Hz and sampled at 1000 Hz during recording. Eye movements and blinks were captured by two pairs of electrooculogram (EOG) electrodes positioned horizontally and vertically around the eyes, respectively. The head position was continuously monitored using 5 head position indicator (HPI) coils attached on the left, middle and right forehead and mastoids. Three anatomical landmarks (the left and right preauricular points and the nasion) and several dozens of extra points around the head surface, along with the HPI coils positions, were digitized for subsequent co-registration of each individual participant’s MEG data with the structural magnetic resonance image (MRI) of their brain.

Structural MRIs were scanned at the ANI Advanced Magnetic Imaging Centre after the MEG session using a 3 T MRI scanner (Magnetom Skyra, Siemens) with a 32-channel head coil and T1-weighted MPRAGE and T2-weighted SPC SAG sequences.

### 2.5 MEG preprocessing

MEG data were analyzed using the MNE-Python software package (Gramfort et al., 2013). MEG sensors with evident noisy signals were detected visually and excluded from analysis. Spatiotemporal signal space separation (tSSS) was applied to remove external environmental noise and compensate for head movements (Taulu & Simola, 2006). Thereafter, the data was band-pass filtered at 0.1–40 Hz. Artifacts associated with eye movements, eye blinks and heartbeats were removed using independent component analysis (ICA). ICA decomposition was estimated on the data additionally high-pass filtered at 1 Hz to approach the ICA’s stationarity assumption (Jas et al., 2018). Artifact-related components initially automatically detected, and following a visual inspection were excluded from the data. Thereafter, we extracted epochs from –200 to 1100 ms with respect to each stimulus presentation, including a –200–0 ms pre-stimulus baseline. Epochs with excessive peakto-peak signal amplitudes were removed, with the cutoff threshold of 3000 fT/cm for the gradiometer sensors and 4000 T for the magnetometer sensors. We then averaged the epochs separately per each condition (minimum 130 epochs).

### 2.6 Analysis of MEG evoked activity

To obtain a preliminary overview of the observed data, we aggregated grand-averaged areal evoked-responses from 204 gradiometers across eight regions: bilateral frontal, temporal, parietal, and occipital areas. Each time course was characterized by calculating the root mean square of the amplitude across areal sensors. To ensure consistency in relative sensor locations across subjects, head positions were aligned by transforming head positions of all subjects into a reference position, based on an participant with an average brain size and head-helmet distance, using Elekta Neuromag MaxFilter software.

We estimated source-level evoked activity using MNE-Python (Gramfort et al., 2013). Participants’ cortical surfaces were first reconstructed from their structural T1 MP-RAGE and T2 SPC SAG images using the Freesurfer software package (Dale, Fischl, & Sereno, 1999; Fischl, Liu, & Dale, 2001; Fischl, Sereno, & Dale, 1999). The minimum-norm estimation method (Hämäläinen & Sarvas, 1989) was then used to estimate the sources of the averaged evoked responses in the four conditions on each individual’s reconstructed cortical surface. A boundary-element model (BEM) was created by stripping the outer skull and scalp from the pial surface using the watershed algorithm in FreeSurfer. A single-layer BEM with an icosahedral mesh of 2,562 vertices per hemisphere was used as a head conductor model in the forward computation. To calculate the inverse operator in each participant, we applied a loose constraint parameter (0.2) to the relative weighting of tangential versus radial current dipole orientations, and a depth weighting parameter (0.8) to to increase the contributions of deeper sources. We constructed and regularized an empirical noise-covariance matrix using the baseline interval of all epochs and a regularization factor of 0.1 for noise-normalized dynamic statistical parametric maps (dSPM; Dale et al., 2000).

For group-level analyses, the individual source estimates were morphed to a standard template brain provided by FreeSurfer (fsaverage).

### 2.7 Regions of interest

We used a custom-made parcellation including 69 and 70 brain regions for left and right hemispheres, respectively, based on the Destrieux Atlas for fsaverage (Ala-Salomäki, Kujala, Liljeström, & Salmelin, 2021). We selected ROIs that were found to be highly and significantly sensitive to misspelling in the present study and also located within canonical language areas identified in prior neuroimaging studies of word reading (Kaestner et al., 2021, 2022; Price & Devlin, 2011; Wydell et al., 2003): left ventral occipitotemporal cortex (vOT), superior temporal cortex (ST; middle part), and precentral cortex (pC; inferior part), as shown in Figure 5b. For completeness, we also examined task effects in their right-hemisphere counterparts.

### 2.8 Statistical testing and modeling

The behavioral results, including accuracy and reaction time, were evaluated with one-way repeated measures analysis of variance (ANOVA) to examine differences in task performance among conditions. Subsequently, post-hoc pairwise *t*-tests were used to identify how the behavioral performance varied as the levels of misspelling increased. The obtained *p* values were corrected using Benjamini–Hochberg false detection rate (FDR) method.

We conducted a one-way repeated measures ANOVA test at the sensor level to obtain the temporal regions that showed significant misspelling sensitivity, which was the basis for selecting time windows of interest. This test was applied on 30-ms, non-overlapping windows. Significance was accepted at a threshold of *p <* 0.01. For the source-level analysis of MEG data across conditions, we employed cluster-based permutation tests to determine significant differences between real and misspelled words across participants. Since this test is not capable of inferring the (spatial or temporal) extents or locations of effects (Sassenhagen & Draschkow, 2019), we performed the test in two ways: First, for addressing the temporal effects we applied the test to the whole cortex in 200-ms time windows. Second, to examine significantly misspelling-sensitive regions, we applied the test to the whole range of time windows between all pairs of conditions in each parcel. In both cases, we used 1024 permutations and a cluster-forming threshold of *t >* 3 based on a one-sample *t*-test. The resulting *p*-values were then corrected for multiple comparisons (across parcels or time), using the Benjamini–Hochberg FDR method. For the analysis of statistical differences between the activation time courses of real and misspelled words within each ROI, we also used a cluster-based permutation test. All parameters were kept the same except for the cluster-forming threshold, which was lowered to *t >* 1.5 based on a one-sample *t*-test to better capture subtle differences. To examine how activation strengths in each ROI differed across conditions in different time windows, we performed one-way repeated measures ANOVA in 200-ms time windows, followed by post-hoc pairwise *t*-tests with Benjamini–Hochberg FDR corrected *p* values.

To further examine the nature of the relationship between brain activation and misspelling levels, we modeled the evoked activity in each ROI using linear mixed effects (LME) analysis (Pinheiro & Bates, 2000). In the LME model, participants were modeled as random effects (random intercepts, fixed slopes), and the fixed effect was the number of replaced letters. We additionally examined the fixed effects of visual distance (visual) and bigram frequency difference between stimulus and its base word (sublexical), as well as stimulus recognizability (lexical). Visual distance was measured by the Euclidean distance between the corresponding pixels of the two rendered images of stimulus and its base word. Recognizability was estimated by the accuracy for each condition obtained from behavioral data. Visual distance and bigram frequency differences were converted into discrete predictor variables by dividing them into four ordinal bins. These bins were designed to ensure that the number of epochs in each bin was nearly equal and sufficient for each participant, approximating the number of epochs in each level of misspelling (∼100–150 epochs per bin depending on how the epochs are distributed in the four bins). The values within each bin were then averaged, and the resulting mean predictors for the four bins were scaled using min-max normalization. An LME model was employed to assess the effects of these predictors on the time course of the evoked responses using 20 ms windows without overlap. The *p*-values obtained from LME at each time point were FDR corrected, with the threshold at *p <* 0.01 for an effect to be considered significant.

## 3 Results

### 3.1 Effect of misspelling on semantic retrieval

To assess how different degrees of misspelling affect semantic retrieval, we examined the behavioral task performance during catch trials. Task accuracy showed a decreasing trend with an increasing number of replaced letters: RW (mean±SD 86.4 ± 16.7%), RL1 (85.2 ± 14.6%), RL2 (67.8 ± 16.0%), and RL3 (49.9 ± 13.0%) (Figure 2). The accuracy dropped significantly from RL1 to RL3 (one-way repeated measures ANOVA: *F* (3, 66) = 60.23, *p <* .001; pairwise *t*-tests across RL1, RL2 and RL3: *t*(22) = 5.81 to 12.00, *p <* .001), with RL3 being not significantly above the chance level accuracy of 50% (*t*(22) = −.05, *p* = .52). However, no significant difference was observed between RW and RL1 (*t*(22) = −.31, *p* = .75). The reaction times increased with an increasing number of replaced letters: RW (1220 ± 226 ms), RL1 (1271 ± 262 ms), RL2 (1431 ± 220 ms), and RL3 (1502 ± 341 ms) (*F* (3, 66) = 15.49, *p <* .001; pairwise *t*-tests across RW, RL1 and RL2: *t*(22) = −7.90 to −2.21, *p <* .001 to .05), plateauing at 2-letter replacement (RL2 vs RL3: *t*(22) = −1.25, *p* = .22).

**Figure 2:**
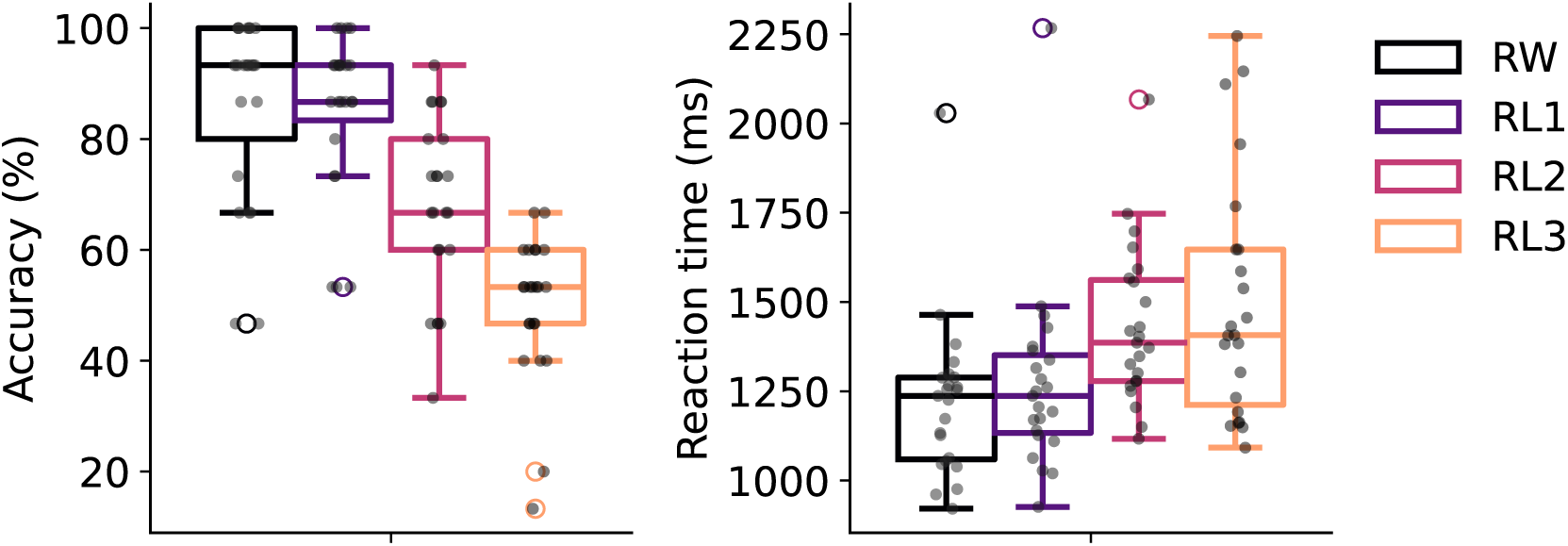
Barplots of task accuracy and reaction time for the four stimulus categories. Each black dot indicates an individual participant’s result(*n* = 23).

### 3.2 Spatiotemporal differences between real and misspelled words in evoked activity

To determine when misspelling sensitivity was reflected in neural responses, we analyzed the grand-averaged areal evoked time courses at the sensor level. As illustrated in Figure 3, we observed that misspelling sensitivity emerged after ∼300 ms in most areas, where misspelled words elicited stronger and more sustained responses than real words. While the initial sensor-level analysis suggested a significant difference between conditions also at ∼200–300 ms in the right occipital area, this effect was not confirmed in the source-level analysis (Figure 3, bottom right). Therefore, in the following, we focus on time windows *>*300 ms.

**Figure 3:**
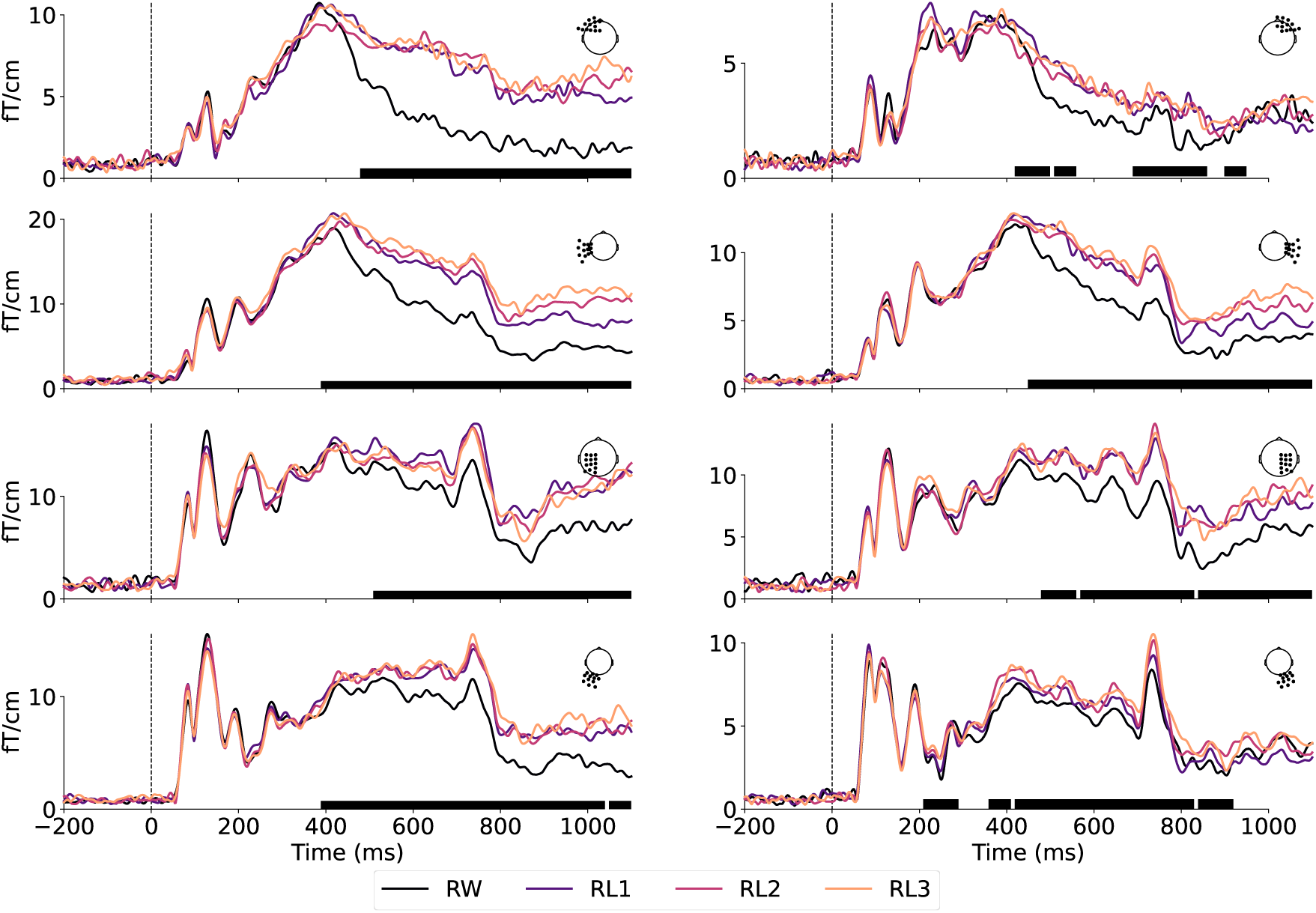
Grand-averaged areal time course of MEG evoked responses across gradiometer sensors located over the frontal (first row), temporal (second row), parietal (third row), and occipital (fourth row) cortex in the left (left column) and right hemisphere (right column). Black bars under each plot indicate regions of significance between all conditions (one-way repeated measures ANOVA, *p <* 0.01).

To investigate the temporally evolving map of misspelling effects at the source level, we contrasted group-level source activation patterns between the three levels of misspelled words and real words across four time windows: 300–500 ms, 500–700 ms, 700–900 ms, and 900–1100 ms (Figure 4a). In the 300–500 ms time window, RL3 elicited stronger activation than RW in the left ST and pC (Figure 4a, row 1). From 500 ms onwards, RL1 and RL2 also produced stronger activity than RW in approximately the same areas, and the right hemisphere was additionally highlighted (Figure 4a, rows 2–3). Furthermore, the left vOT cortex that showed an early response to all stimuli (*<*200 ms; see Supplementary Figure 1 for the group-level evoked responses for each condition separately) was more active for misspelled words than RW from 500 ms onwards. These effects, though diminished over time, sustained until 1100 ms, particularly salient in the left hemisphere.

**Figure 4:**
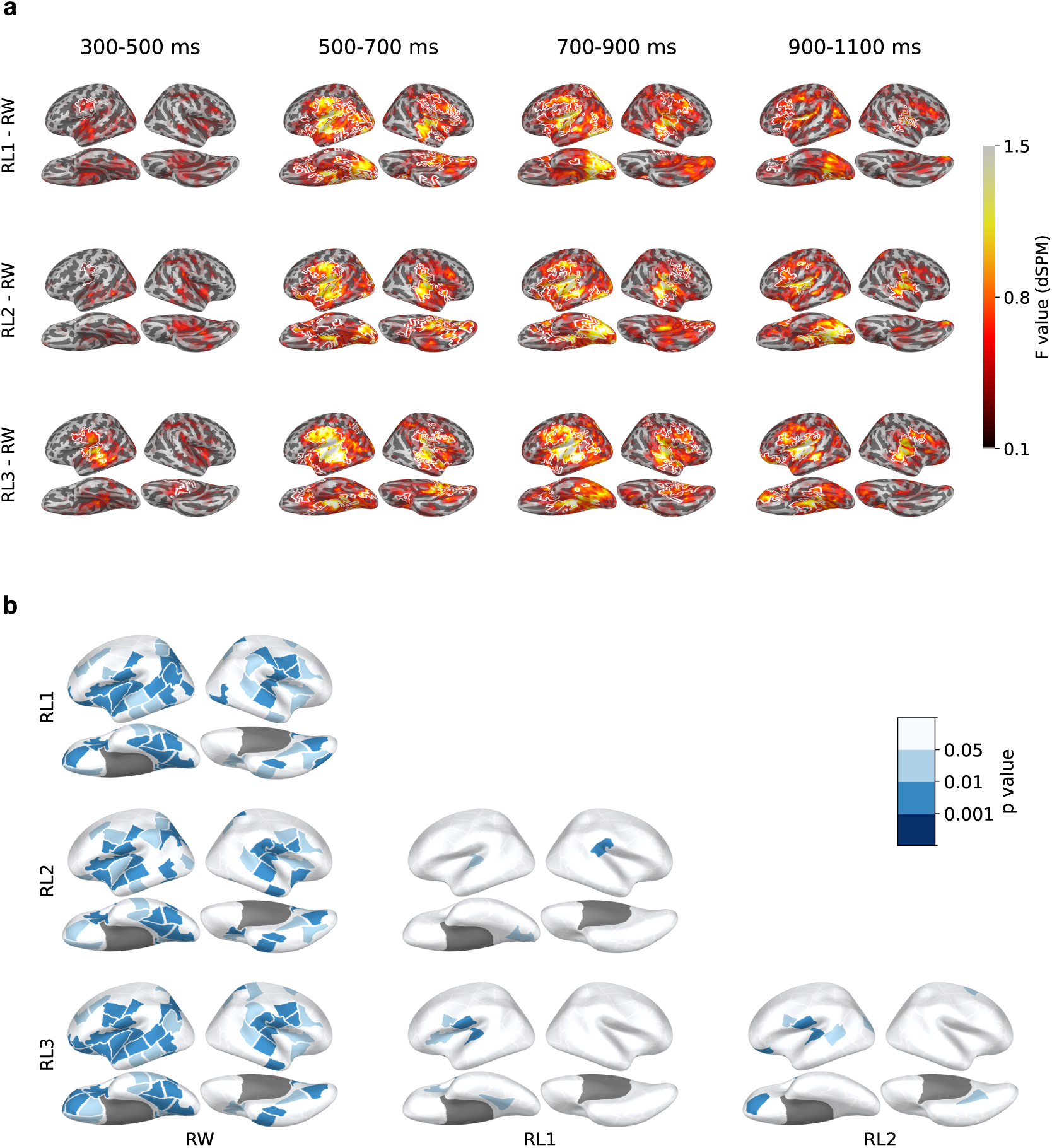
Cortical activation evoked by misspelled words vs. real words during 300–1100 ms. **a**, Group-level source estimates (MNE-dSPM) contrasting RWs and misspelled words in four selected time windows. White borders indicate clusters with *p <* 0.05 in a one-tailed cluster-based permutation test. **b**, Statistical tests on the evoked activity during 300–1100 ms between all pairs of conditions, with FDR-corrected p-values (dark blue, *p <* 0.001; mid blue, *p <* 0.01, light blue, *p <* 0.05, white, *n.s.*).

To examine where in the brain was sensitive to misspellings, we contrasted activation strengths within cortical parcels during the full time window of 300–1100 ms. Significant differences between misspelled and real words were revealed extensively around the perisylvian language regions, predominantly in the left hemisphere (Figure 4b). An increasing number of parcels with a significant effect was observed with increasing levels of misspellings, suggesting more robust neural effects. However, a far smaller set of cortical regions showed significant sensitivity between the increasing levels of misspellings. Nonetheless, stronger activation in left ST and pC was retained for almost all pairs of misspelled words, except that left pC appeared in the contrast between RL1 and RL2. No significant difference between levels of misspelling was found in vOT.

### 3.3 Time courses of activation in the ROIs

To characterize the timing of distinctions between stimulus classes, we focused on three left-hemisphere ROIs: vOT, ST, and pC(Figure 5a). In all ROIs, the activation evoked by the misspelled words differed significantly from that evoked by real words from about 500 ms onwards. We further divided the time courses of activation into discrete time windows and performed pairwise comparisons between conditions (Figure 5b).

**Figure 5:**
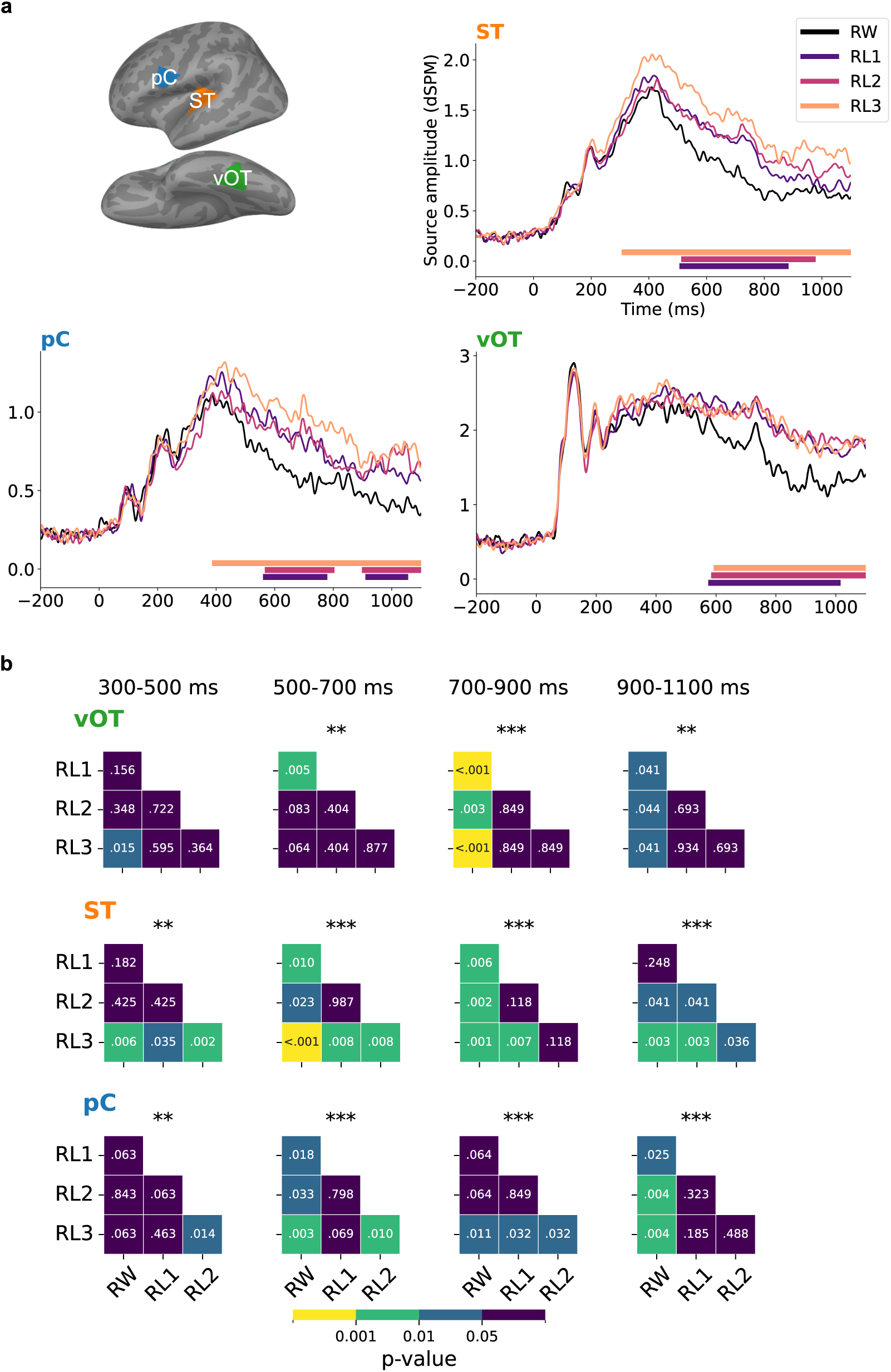
**a**, Regions of interest (ROIs) and averaged evoked responses of each category within the ROIs. Solid bars under the plots indicate the time clusters with *p <* 0.05 based on cluster-based permutation tests between misspelled words and real words. **b**, Results of one-way repeated measures ANOVA and pairwise *t*-test (FDR corrected) between conditions in different time windows. Asterisks above each heatmap indicate the significance level obtained from ANOVA (***, *p <* 0.001; **, *p <* 0.01; *, *p <* 0.05).

The time courses in the vOT exhibited pronounced sensitivity to misspelled words from about 550 ms onwards. Pairwise comparisons revealed that from 700 ms onwards, vOT differentiated all misspelled word types similarly from the real words. Additionally, vOT seemed to show marginally more activation for RL3 than RW at 300-500 ms and for RL1 than RW at 500-700 ms. More anteriorly, ST showed a graded response to increasing misspellings from about 300 ms onwards, a pattern mirrored in pC from about 400 ms onwards. In both ST and pC, the observed activation clusters for RL3 (bars below the time courses) lasted longer and started earlier than those for RL2 and RL1. The pairwise test confirmed and complemented the qualitative observations in ST: RL3 showed task sensitivity in the 300-500 ms time window, thus earlier than the other conditions. Subsequently, RL1 and RL2 also exhibited misspelling effects until the last time window, at which point there was no significant difference between RL1 and RW. In pC, the misspelling effects differed slightly from those in ST and were not as robust. The analysis of corresponding ROIs in the right hemisphere showed notable effects between misspelled and real words primarily in the ST and pC and only few effects in the vOT. Differences between misspelled words were only detectable in the late 900-1100 ms window in the ST (Supplementary Figure 2).

To quantify the relative sensitivity of each ROI to misspelling, we performed a linear mixed effects (LME) analysis with fixed effects of the number of replaced letters (0–3) at each time point during the 0-1100 ms period (Figure 6a). The model over time showed a strong and long-lasting effect that emerged late in the ST at around 350 ms, with a subsequent weaker effect in the pC. A late effect was also observed more posteriorly in the vOT, peaking at around 850 ms.

**Figure 6:**
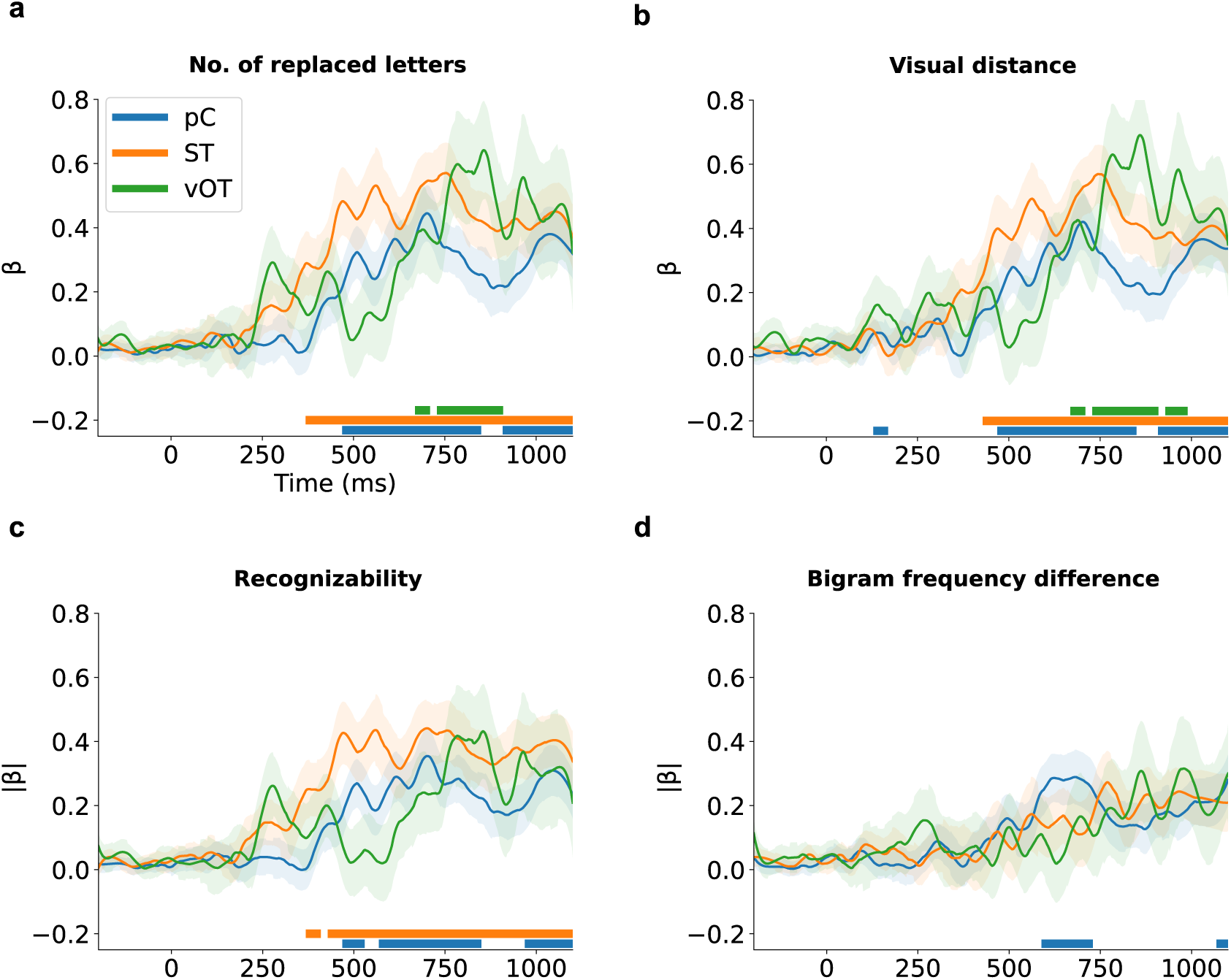
Time courses (LME, *β* ± *s.e.*) of misspelling sensitivity in terms of number of replaced letters. (**a**), visual distance (**b**), bigram frequency difference (**c**) and recognizability (**d**). Solid bars indicate time regions of significant effect (*p <* 0.01).

While the number of manipulated letters was the focus of our study, it is not the only variable influencing the activation patterns of misspelled word reading. Therefore, we also generated LME models to evaluate the effects of additional parameters closely related to misspelling: visual distance, the difference of bigram frequency, and recognizability (Figure 6b-d). We found significant effects of visual distance(Figure 6b), which paralleled those of the number of replaced letters, indicating a correlation between the two variables (Pearson correlation, *r*(538) = 0.95, *p <* 0.001). The pattern of the effects for recognizability appeared similar to that for the number of replaced letters, but no effect was found in the vOT(Figure 6c). Sensitivity to the difference in bigram frequency was observed solely in the pC (Figure 6d).

## 4 Discussion

We aimed to explore where and when the brain activity of skilled readers was modified by the possibility of comprehending misspelled words. Thus, we investigated how increasing degrees of misspelling affect meaning retrieval and cortical dynamics by recording behavioral data and MEG during a silent single-word reading paradigm. The stimuli of misspelled words were perceived to range from real words to complete pseudowords. Our work reveals that left vOT shows selectivity to misspelled words vs. real words, while left ST and pC also differentiate between degrees of misspelling. Notably, these effects occurred late (*>* 300 ms after stimulus onset) and sustained.

### 4.1 Semantic retrieval can survive misspelling

Behavioral data indicated that semantic information could be retrieved to varying degrees from the three categories of misspelled words. RL1 showed no significant differences in accuracy compared to RW, while for RL3 the accuracy plummeted, and RL2 fell between RL1 and RL3, indicating the words had become ambiguous. Therefore, RL1, RL2, and RL3 may be regarded as recognizable, ambiguous, and unrecognizable misspelled words, respectively. The reduced accuracy with an increasing number of misspelled letters was accompanied by a longer reaction time, likely due to deeper rumination required to search for the intended words in the mental lexicon under more challenging conditions. Unlike the masked priming paradigm used in previous behavioral studies of misspelling (Forster et al., 1987; Grainger, 2008; Lupker & Davis, 2009), the present paradigm provided a direct estimate of how well the meaning of misspelled words could be retrieved along the word–pseudoword spectrum.

### 4.2 Left vOT, ST and pC are central cortical regions for disambiguating misspelled words

Salient misspelling effects were observed in left vOT, ST, and pC, which are all regions belonging to the established language network. Hence, it appears that disambiguating misspelled words is handled by this network as well. Broadly, left vOT differentiated the dichotomy of misspelled vs. real words at a coarse level, while left ST and pC distinguished varying levels of misspelling with greater precision.

Left vOT has been linked to prelexical neural processing of letter strings in electrophysiological (MEG, EEG) (Baker et al., 2007; Glezer et al., 2009; Tarkiainen et al., 1999; Vartiainen et al., 2011) and fMRI recordings (Dehaene & Cohen, 2007; Dehaene et al., 2002). We observed notable selectivity for misspelled words in left vOT starting only from 500 ms, with higher activation for pseudowords than words (RW*<*RL3), but no distinction between degrees of misspelling (RL1=RL2=RL3). In MEG studies, left vOT shows a response peaking at around 150–200 ms that is sensitive to letter strings, in general, and thought to reflect pre-lexical processing Tarkiainen et al. (1999). Our results support this view: left vOT showed no early effects on recognizability, which supports the notion that during the first 200 ms, vOT may be involved in processing the visual shape of letters and letter strings, but not in detecting complete word-like forms. However, it is important to note that semantic processing may begin around 250 ms, or even earlier Amsel, Urbach, and Kutas (2013); Hauk, Coutout, Holden, and Chen (2012). This suggests that vOT likely acts as an interface between orthographic and lexico-semantic information. In EEG studies, N250 peaking at around 200–300 ms was found to be sensitive to misspellings in masked priming studies (Gutíerrez-Sigut et al., 2019; Ktori et al., 2014; Meade, Grainger, & Holcomb, 2021; Meade et al., 2020; Meade, Mahnich, et al., 2021). We did not find such an effect in our results. This is probably because in the aforementioned EEG studies, it was the prime that was the misspelled word, but the response to the target word (correctly spelled) was analyzed. The fact that we do not see an N250 effect when we study misspelled words in isolation, in line with a previous EEG study using isolated words (Vergara-Martínez et al., 2013), suggests that during the masked priming studies, the resolution of the misspelled prime had already occurred before the onset of the target word. However, fMRI studies do show involvement of vOT in the resolution of misspelled words (Carreiras et al., 2007; Cattinelli et al., 2013; Fiebach et al., 2002; Heim et al., 2005; Kronbichler et al., 2007; McNorgan et al., 2015), suggesting that they may have picked up the late effect (RL*>*RW after 500 ms) we observed in this study.

While vOT showed no distinction between RL1 and RL2, a more refined distinction between words, misspelled words and pseudowords emerged in left ST and pC. Previous MEG studies have typically linked ST activation in the N400 time window, with lexical-semantic processing during word reading (Helenius, Salmelin, Service, & Connolly, 1998; Salmelin, Schnitzler, Schmitz, & Freund, 2000; Van Petten & Luka, 2006). In studies using (correctly spelled) written words, the activity in ST has been found to be modulated by factors such as task (Chen, Davis, Pulvermüller, & Hauk, 2013), word frequency (Simos et al., 2009), and context (Helenius et al., 1998). In addition, this modulation also extends to word-likeness, with stronger activation for pseudowords than real words (Mainy et al., 2008; Vartiainen et al., 2011; Wydell et al., 2003). In our results, the initial stages of the typical N400 window (300–500 ms) only showed a distinction between pseudowords (RL3) and real words (RW=RL1=RL2) but, thereafter (∼500 ms onward), left ST became sensitive to degrees of misspellings (RW*<*RL1=RL2*<*RL3). This finding was in line with earlier MEG observations on pseudowords compared to real words (Mainy et al., 2008; Wydell et al., 2003). However, the sensitivity to RL1 emerged later than that in a previous EEG study reporting stronger amplitudes for RL1 than RW in the N400 window already (Vergara-Martínez et al., 2013).

Activation patterns in left pC replicated and extended previous fMRI studies that have demonstrated a preference for pseudowords in left frontal operculum, which comprises the inferior portion of left pC (Carreiras et al., 2007; Cattinelli et al., 2013; Fiebach et al., 2002; Heim et al., 2005; Konstantopoulos & Giakoumettis, 2023; Kronbichler et al., 2007; McNorgan et al., 2015). pC has been proposed to be engaged in articulatory phonological processing (Tourville & Guenther, 2011). For example, pC has demonstrated selectivity to lexicality in a study of reading aloud (Woolnough et al., 2022). Additionally, Chen et al. (2013) found more activation in pC for silent reading compared to lexical decision, indicating that silent reading could involve covert articulation. Several silent reading studies have suggested that pC may contribute to the grapheme-to-phoneme conversion process (Kaestner et al., 2021, 2022). The triangle model states that skilled readers can employ two “routes” for recognizing words: For common words, the visual shape is immediately recognized and the appropriate lexical item is activated. For complex or uncommon words, there is a slower phonological route where letters/syllables are mentally sounded out before activation of candidate lexical items (Jobard, Crivello, & Tzourio-Mazoyer, 2003). The heightened involvement of pC during processing of misspelled words may, therefore, indicate that the phonological route is being employed when faced with misspelled words. This interpretation is further supported by the observation that pC demonstrated an exclusive effect of bigram frequency (Figure 6d). It is worth noting that pC does not typically appear as a notably active cortical area in electrophysiological studies of silent single-word reading. The large proportion of misspelled words in the present MEG study may have enhanced phonological processing of all stimuli and, thereby, highlighted the role of pC.

### 4.3 The process for disambiguating misspelled words is late and sustained

Overall, our study shows that when words are presented in isolation, the effect of misspellings only manifests late into the time course of the evoked response, starting from the N400 window and extending for a prolonged time after. This suggests that whatever process is responsible for the resolution of the misspellings, it happens after the initial activation of candidates in the lexicon.

We could be seeing the effects of increased lexico-semantic prediction error, as suggested by the latest predictive coding model of the N400 (Eddine et al., 2024). According to this theory, the bottom-up visual input activates multiple lexical candidates that are orthographically similar, which are then refined through a top-down prediction process that reconciles the activated lexical items with the original input. When a word is correctly spelled (RW), this reconciliation can happen quickly and unambiguously, as there is only a single lexical item that perfectly matches the input. However, in the case of misspelled words and pseudowords, the input can never be fully reconciled with the activated lexical items, which shows as a sustained prediction error. This may also explain why, in vOT, we first see a distinction between pseudowords (RL3) and correctly spelled words (RW), and only later a distinction between degrees of misspellings (RL1 and RL2 versus RW) (Figure 5b). If we assume that RW, RL1 and RL2 initially activate the same lexical items, while RL3 activates a wider selection of items, RW/R1/R2 would initially produce roughly the same amount of prediction error, while RL3 produces more.

Our results indicate that the initial buildup of lexico-semantic activation followed by its gradual decay is a process occurring within the language network, with special roles for vOT, ST and pC. The particularly late effect in vOT (after 500 ms), an area typically associated with “low-level” pre-lexical processing, points to an anterior-to-posterior spread of misspelling information, potentially in a top-down manner from left ST/pC to vOT (Figure 6). In this manner, the reading of misspelled words seems to align with predictive coding theory (Friston, 2010). Specifcally, the comparable activation strengths in left vOT for all misspelled conditions (RL1 = RL2 = RL3 *>* RW) suggest uniform prediction errors at the lexico-semantic level, where the orthographic representations fail to match any lexical entry equally for all levels of misspelled words (Price & Devlin, 2011; Woolnough et al., 2021). However, the activation strengths are graded in left ST and pC, reflecting less prediction errors for recognizable misspelled words than unrecognizable ones based on top-down predictions from prior knowledges.

This result contrasts with the rapid feedforward model of word recognition, which is typically completed within 500 ms (Pammer et al., 2004). Instead, similar recurrent feedforward–feedback processing mechanism has also been suggested in several visual object recognition studies (Karimi-Rouzbahani, Ramezani, Woolgar, Rich, & Ghodrati, 2021; Kietzmann et al., 2019; Rajaei, Mohsenzadeh, Ebrahimpour, & Khaligh-Razavi, 2019; Von Seth, Nicholls, Tyler, & Clarke, 2023). We postulate that misspelled word reading may provide circumstances under which the brain would choose to wait for additional top-down constraints (Carreiras, Armstrong, Perea, & Frost, 2014). Future work needs to evaluate theories of top-down processing (such as the interactive account for vOT’s function (Price & Devlin, 2011) and predictive coding theory for N400 component (Eddine et al., 2024)) by investigating how regions interact to process and transform information (Hauk, Jackson, & Rahimi, 2023).

## 5 Conclusion

This study delineated the cortical dynamics during silent misspelled word reading. By systematically manipulating the degrees of misspelling, we generated a stimulus spectrum ranging from recognizable to unrecognizable misspelled words, confirmed by the behavioral results. Left vOT, ST, and pC, typically associated with orthographic, lexico-semantic, and phonological processing, respectively, were engaged in a late and sustained process to disambiguate misspelled words from about 300 ms onwards. These results seem to conflict with the general concept of rapid feedforward process of word recognition. The remarkably late effect of misspelling in left vOT speaks to an anterior-to-posterior spread of misspelling information. Such potential feedback and feedforward interactions need to be validated by future studies.

## 6 Data availability statement

The stimulus list (including Presentation scripts) and MEG derivatives necessary for reproducing the results in this study are available at https://osf.io/r6e7z. In accordance with the EU General Data Protection Regulation that limits sharing of personal data, only MEG data that have been grand-averaged or morphed to a template brain are provided. The cortical parcellations used in this study are available at https://github.com/ AaltoImagingLanguage/custom parcellations.

## 7 Code availability statement

For preprocessing we used MNE-Python (Gramfort et al., 2013), Maxfilter, and FreeSurfer software. For LME model, we used the statsmodels package (Seabold & Perktold, 2010) in Python. For other analysis, we used the following packages: MNE-Python (Gramfort et al., 2013), pingouin (Vallat, 2018), NumPy (Harris et al., 2020), Pandas (Wes McKinney, 2010), and Matplotlib (Hunter, 2007). The custom code used in the study can be accessed at https://github.com/AaltoImagingLanguage/you2024

## 8 Acknowledgments

We would like to thank Olaf Hauk for his valuable comments on the manuscript. Thanks to Mia Illman and Rami Kunnas for their assistance with data collection. We also acknowledge the computational resources provided by the Aalto Science-IT project. This work was funded by the Academy of Finland (#346585 and #343385 to M.v.V., #355407 to R.S.), the China Scholarship Council (#202206100022 to J.Y.) and the Sigrid Juśelius Foundation (to R.S.).

## 9 Author contributions

Conceptualization: J.Y., M.v.V., R.S.; Data collection: J.Y., A.S.; Methodology: J.Y., M.v.V., T.L-K, A.S., R.S.; Interpretation: J.Y., M.v.V., R.S.; Writing—original draft, J.Y.; Writing—review & editing, J.Y., M.v.V., T.L-K, A.S., R.S.

## References

1. Ala-Salomäki, H., Kujala, J., Liljeström, M., & Salmelin, R. (2021). Picture naming yields highly consistent cortical activation patterns: Test–retest reliability of magnetoencephalography recordings. NeuroImage, 227, 117651.

2. Almeida, D., & Poeppel, D. (2013). Word-specific repetition effects revealed by meg and the implications for lexical access. Brain and language, 127 (3), 497–509.

3. Amsel, B. D., Urbach, T. P., & Kutas, M. (2013). Alive and grasping: Stable and rapid semantic access to an object category but not object graspability. NeuroImage, 77, 1–13.

4. Baker, C. I., Liu, J., Wald, L. L., Kwong, K. K., Benner, T., & Kanwisher, N. (2007). Visual word processing and experiential origins of functional selectivity in human extrastriate cortex. Proceedings of the National Academy of Sciences, 104 (21), 9087–9092.

5. Carreiras, M., Armstrong, B. C., Perea, M., & Frost, R. (2014). The what, when, where, and how of visual word recognition. Trends in cognitive sciences, 18 (2), 90–98.

6. Carreiras, M., Mechelli, A., Estévez, A., & Price, C. J. (2007). Brain activation for lexical decision and reading aloud: two sides of the same coin? Journal of cognitive neuroscience, 19 (3), 433–444.

7. Cattinelli, I., Borghese, N. A., Gallucci, M., & Paulesu, E. (2013). Reading the reading brain: a new meta-analysis of functional imaging data on reading. Journal of neurolinguistics, 26 (1), 214–238.

8. Chen, Y., Davis, M. H., Pulvermüller, F., & Hauk, O. (2013). Task modulation of brain responses in visual word recognition as studied using eeg/meg and fmri. Frontiers in Human Neuroscience, 7, 376.

9. Chen, Y., Davis, M. H., Pulvermüller, F., & Hauk, O. (2015). Early visual word processing is flexible: Evidence from spatiotemporal brain dynamics. Journal of Cognitive Neuroscience, 27 (9), 1738–1751.

10. Dale, A. M., Fischl, B., & Sereno, M. I. (1999). Cortical surface-based analysis: I. segmentation and surface reconstruction. Neuroimage, 9 (2), 179–194.

11. Dale, A. M., Liu, A. K., Fischl, B., Buckner, R. L., Belliveau, J. W., Lewine, J. D., & Halgren, E. (2000). Dynamic statistical parametric mapping: combining fmri and meg for high-resolution imaging of cortical activity. neuron, 26 (1), 55–67.

12. Damerau, F. J. (1964, March). A technique for computer detection and correction of spelling errors. Communications of the ACM, 7 (3), 171–176. Retrieved 2024-0424, from 10.1145/363958.363994 doi: 10.1145/363958.363994

13. Davis, M. (2013). Aoccdrnig to a rscheearch at cmabrigde uinervtisy … according to a research (sic) at cambridge university. Online: Internet http://www.mrccbu.cam.ac.uk/people/matt.davis/Cmabrigde/10thApril.

14. Dehaene, S., & Cohen, L. (2007). Cultural recycling of cortical maps. Neuron, 56 (2), 384–398.

15. Dehaene, S., & Cohen, L. (2011). The unique role of the visual word form area in reading. Trends in cognitive sciences, 15 (6), 254–262.

16. Dehaene, S., Le Clec’H, G., Poline, J.-B., Le Bihan, D., & Cohen, L. (2002). The visual word form area: a prelexical representation of visual words in the fusiform gyrus. Neuroreport, 13 (3), 321–325.

17. Eddine, S. N., Brothers, T., Wang, L., Spratling, M., & Kuperberg, G. R. (2024). A predictive coding model of the n400. Cognition, 246, 105755.

18. Evans, G. A., Lambon Ralph, M. A., & Woollams, A. M. (2012). What’s in a word? a parametric study of semantic influences on visual word recognition. Psychonomic bulletin & review, 19, 325–331.

19. Fiebach, C. J., Friederici, A. D., Müller, K., & Von Cramon, D. Y. (2002). fmri evidence for dual routes to the mental lexicon in visual word recognition. Journal of cognitive neuroscience, 14 (1), 11–23.

20. Fischl, B., Liu, A., & Dale, A. M. (2001). Automated manifold surgery: constructing geometrically accurate and topologically correct models of the human cerebral cortex. IEEE transactions on medical imaging, 20 (1), 70–80.

21. Fischl, B., Sereno, M. I., & Dale, A. M. (1999). Cortical surface-based analysis: Ii: inflation, flattening, and a surface-based coordinate system. Neuroimage, 9 (2), 195–207.

22. Forster, K. I., Davis, C., Schoknecht, C., & Carter, R. (1987). Masked priming with graphemically related forms: Repetition or partial activation? The Quarterly Journal of Experimental Psychology, 39 (2), 211–251.

23. Friston, K. (2010). The free-energy principle: a unified brain theory? Nature reviews neuroscience, 11 (2), 127–138.

24. Glezer, L. S., Jiang, X., & Riesenhuber, M. (2009). Evidence for highly selective neuronal tuning to whole words in the “visual word form area”. Neuron, 62 (2), 199–204.

25. Gow Jr, D. W., Segawa, J. A., Ahlfors, S. P., & Lin, F.-H. (2008). Lexical influences on speech perception: a granger causality analysis of meg and eeg source estimates. Neuroimage, 43 (3), 614–623.

26. Grainger, J. (2008). Cracking the orthographic code: An introduction. Language and cognitive processes, 23 (1), 1–35.

27. Grainger, J., & Jacobs, A. M. (1996). Orthographic processing in visual word recognition: a multiple read-out model. Psychological review, 103 (3), 518.

28. Gramfort, A., Luessi, M., Larson, E., Engemann, D. A., Strohmeier, D., Brodbeck, C., . . . Hämäläinen, M. (2013, December). MEG and EEG data analysis with MNE-Python. Frontiers in Neuroscience, 7 . Retrieved 2024-0424, from https://www.frontiersin.org/journals/neuroscience/articles/10.3389/fnins.2013.00267/full (Publisher: Frontiers) doi: 10.3389/fnins.2013.00267

29. Gutíerrez-Sigut, E., Marcet, A., & Perea, M. (2019). Tracking the time course of letter visual-similarity effects during word recognition: A masked priming erp investigation. Cognitive, Affective, & Behavioral Neuroscience, 19 (4), 966–984.

30. Harris, C. R., Millman, K. J., van der Walt, S. J., Gommers, R., Virtanen, P., Cournapeau, D., . . . Oliphant, T. E. (2020). Array programming with NumPy. Nature, 585, 357–362. doi: 10.1038/s41586-020-2649-2

31. Hauk, O., Coutout, C., Holden, A., & Chen, Y. (2012). The time-course of single-word reading: evidence from fast behavioral and brain responses. NeuroImage, 60 (2), 1462–1477.

32. Hauk, O., Jackson, R. L., & Rahimi, S. (2023). Transforming the neuroscience of language: estimating pattern-to-pattern transformations of brain activity. Language, Cognition and Neuroscience, 1–16.

33. Hauk, O., Pulvermüller, F., Ford, M., Marslen-Wilson, W., & Davis, M. H. (2009). Can i have a quick word? early electrophysiological manifestations of psycholinguistic processes revealed by event-related regression analysis of the eeg. Biological psychology, 80 (1), 64–74.

34. Heim, S., Alter, K., Ischebeck, A. K., Amunts, K., Eickhoff, S. B., Mohlberg, H., . . . Friederici, A. D. (2005). The role of the left brodmann’s areas 44 and 45 in reading words and pseudowords. Cognitive Brain Research, 25 (3), 982–993.

35. Helenius, P., Salmelin, R., Service, E., & Connolly, J. F. (1998). Distinct time courses of word and context comprehension in the left temporal cortex. Brain: a journal of neurology, 121 (6), 1133–1142.

36. Holcomb, P. J., Grainger, J., & O’rourke, T. (2002). An electrophysiological study of the effects of orthographic neighborhood size on printed word perception. Journal of cognitive neuroscience, 14 (6), 938–950.

37. Hultén, A., van Vliet, M., Kivisaari, S., Lammi, L., Lindh-Knuutila, T., Faisal, A., & Salmelin, R. (2021, October). The neural representation of abstract words may arise through grounding word meaning in language itself. Human Brain Mapping, 42 (15), 4973–4984. doi: 10.1002/hbm.25593

38. Hunter, J. D. (2007). Matplotlib: A 2d graphics environment. Computing in Science & Engineering, 9 (3), 90–95. doi: 10.1109/MCSE.2007.55

39. Hämäläinen, M. S., & Sarvas, J. (1989). Realistic conductivity geometry model of the human head for interpretation of neuromagnetic data. IEEE transactions on biomedical engineering, 36 (2), 165–171.

40. Jas, M., Larson, E., Engemann, D. A., Leppäkangas, J., Taulu, S., Hämäläinen, M., & Gramfort, A. (2018, August). A Reproducible MEG/EEG Group Study With the MNE Software: Recommendations, Quality Assessments, and Good Practices. Frontiers in Neuroscience, 12 . Retrieved 2024-0424, from https://www.frontiersin.org/journals/neuroscience/articles/10.3389/fnins.2018.00530/full (Publisher: Frontiers) doi: 10.3389/fnins.2018.00530

41. Jobard, G., Crivello, F., & Tzourio-Mazoyer, N. (2003). Evaluation of the dual route theory of reading: A metanalysis of 35 neuroimaging studies. NeuroImage, 20 (2), 693–712. doi: 10.1016/S1053-8119(03)00343-4

42. Kaestner, E., Thesen, T., Devinsky, O., Doyle, W., Carlson, C., & Halgren, E. (2021). An intracranial electrophysiology study of visual language encoding: the contribution of the precentral gyrus to silent reading. Journal of Cognitive Neuroscience, 33 (11), 2197–2214.

43. Kaestner, E., Wu, X., Friedman, D., Dugan, P., Devinsky, O., Carlson, C., . . . Halgren, E. (2022). The precentral gyrus contributions to the early time-course of grapheme-to-phoneme conversion. Neurobiology of Language, 3 (1), 18–45.

44. Kanerva, J., Luotolahti, J., Laippala, V., & Ginter, F. (2014). Syntactic n-gram collection from a large-scale corpus of internet finnish. In Human language technologies–the baltic perspective (pp. 184–191). IOS Press.

45. Karimi-Rouzbahani, H., Ramezani, F., Woolgar, A., Rich, A., & Ghodrati, M. (2021). Perceptual difficulty modulates the direction of information flow in familiar face recognition. NeuroImage, 233, 117896.

46. Kietzmann, T. C., Spoerer, C. J., Sorensen, L. K., Cichy, R. M., Hauk, O., & Kriegeskorte, N. (2019). Recurrence is required to capture the representational dynamics of the human visual system. Proceedings of the National Academy of Sciences, 116 (43), 21854–21863.

47. Konstantopoulos, K., & Giakoumettis, D. (2023). Chapter 1 – basic knowledge on neuroanatomy and neurophysiology of the central nervous system. In K. Konstantopoulos & D. Giakoumettis (Eds.), Neuroimaging in neurogenic communication disorders (p. 1–30). Academic Press. Retrieved from https://www.sciencedirect.com/science/article/pii/B9780128238752000049 doi: 10.1016/ B978-0-12-823875-2.00004-9

48. Kronbichler, M., Bergmann, J., Hutzler, F., Staffen, W., Mair, A., Ladurner, G., & Wimmer, H. (2007). Taxi vs. taksi: on orthographic word recognition in the left ventral occipitotemporal cortex. Journal of cognitive neuroscience, 19 (10), 1584–1594.

49. Ktori, M., Kingma, B., Hannagan, T., Holcomb, P. J., & Grainger, J. (2014). On the time-course of adjacent and non-adjacent transposed-letter priming. Journal of Cognitive Psychology, 26 (5), 491–505.

50. Kutas, M., & Federmeier, K. D. (2011). Thirty years and counting: finding meaning in the n400 component of the event-related brain potential (erp). Annual review of psychology, 62, 621–647.

51. Levenshtein, V. I. (1965). Binary codes capable of correcting deletions, insertions, and reversals. Soviet physics. Doklady . Retrieved 2024-04-24, from https://www.semanticscholar.org/paper/Binary-codes-capable-of-correcting-deletions%2C-and-Levenshtein/b2f8876482c97e804bb50a5e2433881ae31d0cdd

52. Lupker, S. J., & Davis, C. J. (2009). Sandwich priming: a method for overcoming the limitations of masked priming by reducing lexical competitor effects. *Journal of Experimental Psychology: Learning*, Memory, and Cognition, 35 (3), 618.

53. Mainy, N., Jung, J., Baciu, M., Kahane, P., Schoendorff, B., Minotti, L., . . . Lachaux, J.-P. (2008). Cortical dynamics of word recognition. Human brain mapping, 29 (11), 1215–1230.

54. Martin, R. C., Tan, Y., Newsome, M. R., & Vu, H. (2017). Language and lexical processing. In Reference module in neuroscience and biobehavioral psychology. Elsevier. Retrieved from https://www.sciencedirect.com/science/article/pii/ B9780128093245030789 doi: 10.1016/B978-0-12-809324-5.03078-9

55. McNorgan, C., Chabal, S., O’Young, D., Lukic, S., & Booth, J. R. (2015). Task dependent lexicality effects support interactive models of reading: A meta-analytic neuroimaging review. Neuropsychologia, 67, 148–158.

56. Meade, G., Grainger, J., & Holcomb, P. J. (2021). Language dominance modulates transposed-letter n400 priming effects in bilinguals. Journal of Cognition, 5 (1).

57. Meade, G., Grainger, J., Midgley, K. J., Holcomb, P. J., & Emmorey, K. (2020). An erp investigation of orthographic precision in deaf and hearing readers. Neuropsychologia, 146, 107542.

58. Meade, G., Mahnich, C., Holcomb, P. J., & Grainger, J. (2021). Orthographic neighborhood density modulates the size of transposed-letter priming effects. *Cognitive, Affective*, & Behavioral Neuroscience, 21, 948–959.

59. Pammer, K., Hansen, P. C., Kringelbach, M. L., Holliday, I., Barnes, G., Hillebrand, A., . . . Cornelissen, P. L. (2004). Visual word recognition: the first half second. Neuroimage, 22 (4), 1819–1825.

60. Pearson, P. D., Barr, R., Kamil, M. L., & Mosenthal, P. (1984). Handbook of reading research, volume 1.

61. Pinheiro, J. C., & Bates, D. M. (2000). Linear mixed-effects models: basic concepts and examples. Mixed-effects models in S and S-Plus, 3–56.

62. Price, C. J., & Devlin, J. T. (2011). The interactive account of ventral occipitotemporal contributions to reading. Trends in cognitive sciences, 15 (6), 246–253.

63. Rajaei, K., Mohsenzadeh, Y., Ebrahimpour, R., & Khaligh-Razavi, S.-M. (2019). Beyond core object recognition: Recurrent processes account for object recognition under occlusion. PLoS computational biology, 15 (5), e1007001.

64. Salmelin, R., Kujala, J., & Liljeström, M. (2019). Magnetoencephalography and the cortical dynamics of language processing. Handbook of Neurolinguistics .

65. Salmelin, R., Schnitzler, A., Schmitz, F., & Freund, H.-J. (2000). Single word reading in developmental stutterers and fluent speakers. Brain, 123 (6), 1184–1202.

66. Sassenhagen, J., & Draschkow, D. (2019). Cluster-based permutation tests of meg/eeg data do not establish significance of effect latency or location. Psychophysiology, 56 (6), e13335.

67. Schuster, S., Hawelka, S., Richlan, F., Ludersdorfer, P., & Hutzler, F. (2015). Eyes on words: A fixation-related fmri study of the left occipito-temporal cortex during self-paced silent reading of words and pseudowords. Scientific reports, 5 (1), 12686.

68. Seabold, S., & Perktold, J. (2010). statsmodels: Econometric and statistical modeling with python. In 9th python in science conference.

69. Simos, P. G., Pugh, K., Mencl, E., Frost, S., Fletcher, J. M., Sarkari, S., & Papanicolaou, A. C. (2009). Temporal course of word recognition in skilled readers: A magnetoencephalography study. Behavioural brain research, 197 (1), 45–54.

70. Suomi, K., Toivanen, J., & Ylitalo, R. (2009, January). Finnish sound structure: phonetics, phonology, phonotactics and prosody [book]. Retrieved 2024-04-24, from https://oulurepo.oulu.fi/handle/10024/36099 (Accepted: 2023-05-12T05:27:23Z ISBN: 9789514289842 Journal Abbreviation: Finnish sound structure: Phonetics, phonology, phonotactics and prosody Publisher: University of Oulu)

71. Tarkiainen, A., Helenius, P., Hansen, P. C., Cornelissen, P. L., & Salmelin, R. (1999). Dynamics of letter string perception in the human occipitotemporal cortex. Brain, 122 (11), 2119–2132.

72. Taulu, S., & Simola, J. (2006, April). Spatiotemporal signal space separation method for rejecting nearby interference in MEG measurements. Physics in Medicine and Biology, 51 (7), 1759–1768. doi: 10.1088/0031-9155/51/7/008

73. Taylor, J., Rastle, K., & Davis, M. H. (2013). Can cognitive models explain brain activation during word and pseudoword reading? a meta-analysis of 36 neuroimaging studies. Psychological bulletin, 139 (4), 766.

74. Tourville, J. A., & Guenther, F. H. (2011). The diva model: A neural theory of speech acquisition and production. Language and cognitive processes, 26 (7), 952–981.

75. Vallat, R. (2018). Pingouin: statistics in python. J. Open Source Softw., 3 (31), 1026.

76. Van Petten, C., & Luka, B. J. (2006). Neural localization of semantic context effects in electromagnetic and hemodynamic studies. Brain and language, 97 (3), 279–293.

77. Vartiainen, J., Liljeström, M., Koskinen, M., Renvall, H., & Salmelin, R. (2011). Functional magnetic resonance imaging blood oxygenation level-dependent signal and magnetoencephalography evoked responses yield different neural functionality in reading. Journal of Neuroscience, 31 (3), 1048–1058.

78. Vergara-Martínez, M., Perea, M., Gómez, P., & Swaab, T. Y. (2013). Erp correlates of letter identity and letter position are modulated by lexical frequency. Brain and language, 125 (1), 11–27.

79. Von Seth, J., Nicholls, V. I., Tyler, L. K., & Clarke, A. (2023). Recurrent connectivity supports higher-level visual and semantic object representations in the brain. Communications Biology, 6 (1), 1207.

80. Wes McKinney. (2010). Data Structures for Statistical Computing in Python. In Stéfan van der Walt & Jarrod Millman (Eds.), Proceedings of the 9th Python in Science Conference (p. 56–61). doi: 10.25080/Majora-92bf1922-00a

81. Woolnough, O., Donos, C., Curtis, A., Rollo, P. S., Roccaforte, Z. J., Dehaene, S., . . . Tandon, N. (2022). A spatiotemporal map of reading aloud. Journal of Neuroscience, 42 (27), 5438–5450.

82. Woolnough, O., Donos, C., Rollo, P. S., Forseth, K. J., Lakretz, Y., Crone, N. E., . . . Tandon, N. (2021). Spatiotemporal dynamics of orthographic and lexical processing in the ventral visual pathway. Nature human behaviour, 5 (3), 389–398.

83. Wydell, T. N., Vuorinen, T., Helenius, P., & Salmelin, R. (2003). Neural correlates of letter-string length and lexicality during reading in a regular orthography. Journal of Cognitive Neuroscience, 15 (7), 1052–1062.

