## Supplemental material for "Misspelled-word reading modulates late cortical dynamics"

### Supplementary materials

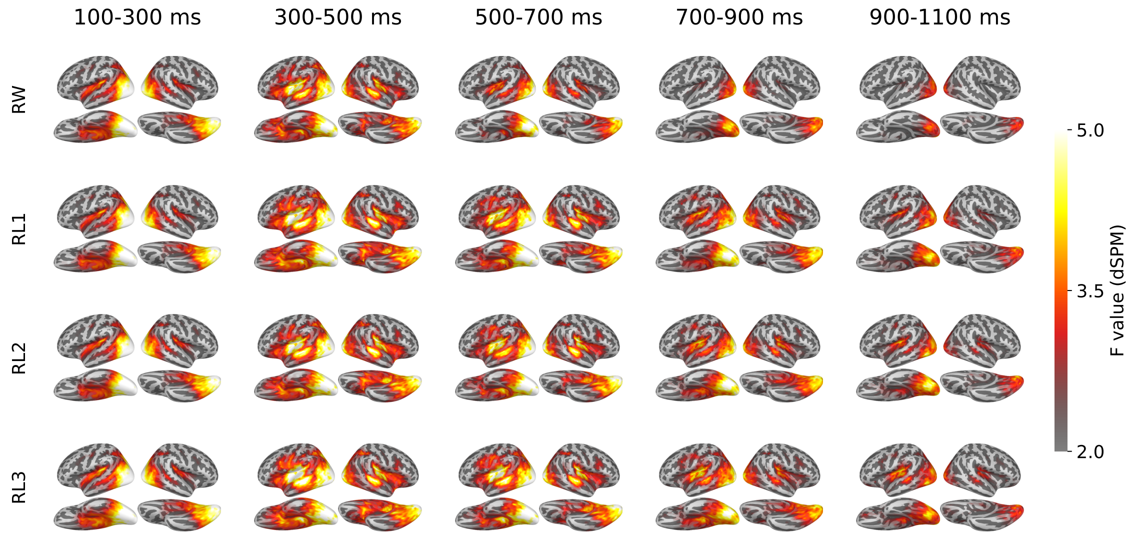


Supplementary Figure 1: Group-level source estimates (MNE-dSPM) for each condition in four selected time windows.

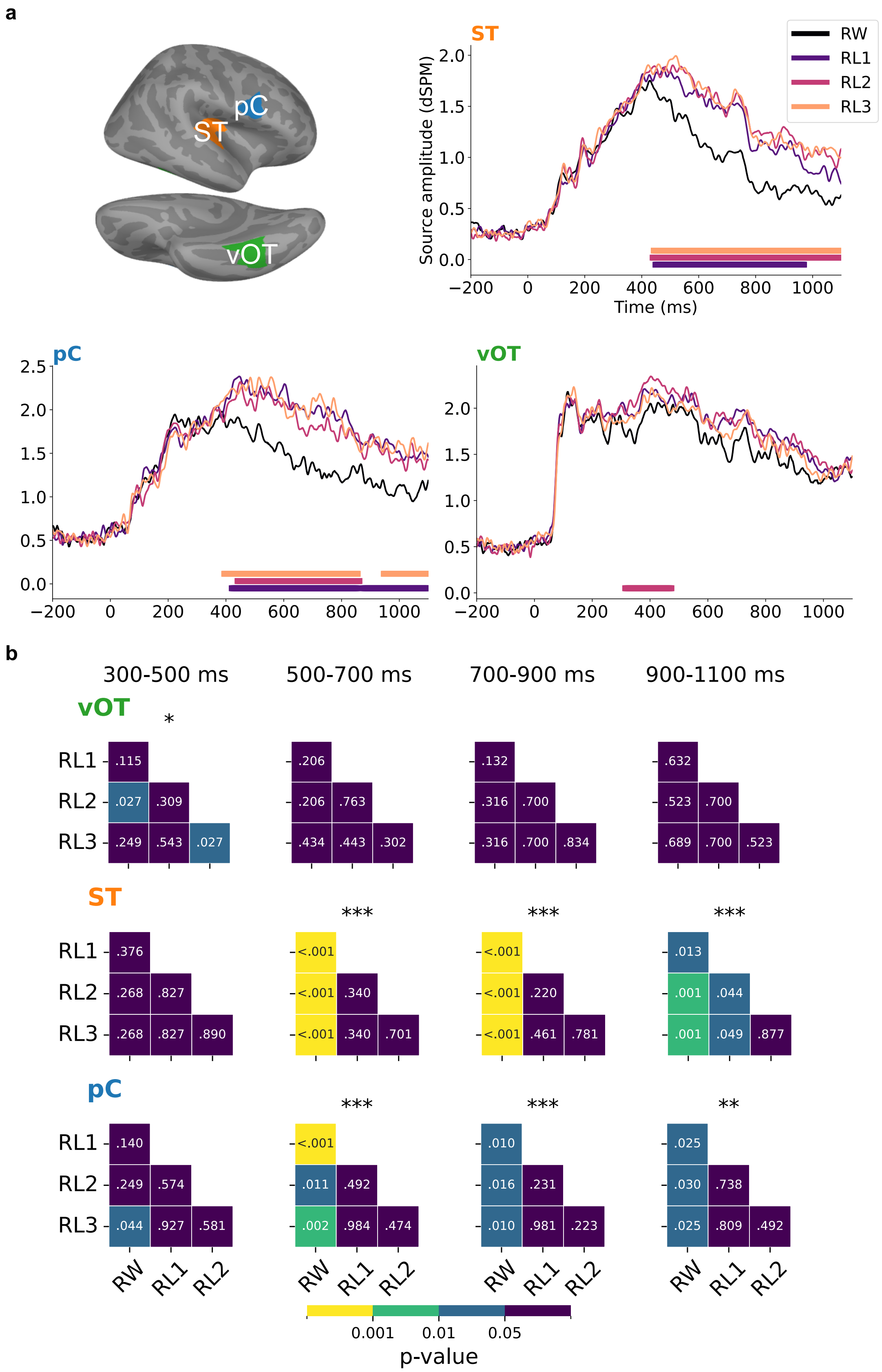


Supplementary Figure 2: **a**, ROIs in the right hemisphere, and averaged evoked responses of each category within the ROIs. Solid bars under the plots indicate the time clusters associated with $p < 0.05$ based on cluster-based permutation tests between misspelled words and real words. **b**, One-way repeated measures ANOVA and Pairwise $t$-test (FDR corrected) between conditions in each time window. Asterisks above each heatmap indicate the significance level obtained from ANOVA (***, $p < 0.001$; **, $p < 0.01$; *, $p < 0.05$).
